# Weaning age and its effect on the development of the swine gut microbiome and resistome

**DOI:** 10.1101/2021.05.20.444551

**Authors:** Devin B Holman, Katherine E. Gzyl, Kathy T. Mou, Heather K. Allen

**Author notes:** Corresponding Author: Devin B. Holman.

## Abstract

Piglets are often weaned between 19 and 22 d of age in North America although in some swine operations this may occur at 14 d or less. Piglets are abruptly separated from their sow at weaning and are quickly transitioned from sow’s milk to a plant-based diet. The effect of weaning age on the long-term development of the pig gut microbiome is largely unknown. Here, pigs were weaned at either 14, 21, or 28 d of age and fecal samples collected 20 times from d 4 (neonatal) through to marketing at d 140. The fecal microbiome was characterized using 16S rRNA gene and shotgun metagenomic sequencing. The fecal microbiome of all piglets shifted significantly three to seven days post-weaning with an increase in microbial diversity. Several *Prevotella* spp. increased in relative abundance immediately after weaning as did butyrate-producing species such as *Butyricicoccus porcorum, Faecalibacterium prausnitzii*, and *Megasphaera elsdenii*. Within 7 days of weaning, the gut microbiome of pigs weaned at 21 and 28 days of age resembled that of pigs weaned at 14 d. Resistance genes to most antimicrobial classes decreased in relative abundance post-weaning with the exception of those conferring resistance to tetracyclines and macrolides-lincosamides-streptogramin B. The relative abundance of microbial carbohydrate-active enzymes (CAZymes) changed significantly in the post-weaning period with an enrichment of CAZymes involved in degradation of plant-derived polysaccharides. These results demonstrate that the pig gut microbiome tends change in a predictable manner post-weaning and that weaning age has only a temporary effect on this microbiome.

**Importance:** Piglets are abruptly separated from their sow at weaning and are quickly transitioned from sow’s milk to a plant-based diet. This is the most important period in commercial swine production yet the effect of weaning age on the long-term development of the pig gut microbiome is largely unknown. Metagenomic sequencing allows for a higher resolution assessment of the pig gut microbiome and enables characterization of the resistome. Here we used metagenomic sequencing to identify bacterial species that were enriched post-weaning and therefore may provide targets for future manipulation studies. In addition, functional profiling of the microbiome indicated that many carbohydrate and metabolic enzymes decrease in relative abundance of after weaning. This study also highlights the challenges faced in reducing antimicrobial resistance in pigs as genes conferring tetracycline and macrolide resistance remained relatively stable from 7 days of age through to market weight at 140 d despite no exposure to antimicrobials.

## Introduction

In commercial swine production, the suckling-weaning transition is the most critical period for piglet health. When piglets are weaned they are abruptly separated from their sow and their diet is changed from an easily digestible milk-based one to a more complex plant-based diet. The risk of developing health problems is increased as piglets are subjected to stress as a result of mixing with unfamiliar piglets, handling, and separation from the sow (1). This stress frequently leads to reduced feed intake immediately following weaning which negatively affects growth performance (2). Consequently, newly weaned piglets frequently develop post-weaning diarrhea, resulting in significant economic losses due to associated piglet morbidity, mortality, and treatment (3). Weaning times vary but piglets can be weaned as young as 14 d or less in some in North American commercial swine operations. Earlier weaning ages allow for a greater number of piglets weaned per sow per year and may also decrease the risk of transmission of certain pathogens from the sow to piglets. However, piglets that are weaned relatively early may be more susceptible to disease and other complications (4).

As with humans and other mammals, the gut microbiome is an important factor affecting swine health. There are an estimated 17 million plus microbial genes in the pig gut microbiome (5) compared to 20,000 to 25,000 genes in the swine genome (6). This greatly expands the genetic potential of the host, particularly as certain microbes can metabolize otherwise non-digestible dietary carbohydrates into a usable energy source. It has been well documented that the pig gut microbiome undergoes a rapid shift following weaning including a decrease in members of the Proteobacteria phylum and *Bacteroides* genus and an increase in genera such as *Prevotella*, *Roseburia*, and *Succinivibrio* (7–10). However, relatively little is known about how weaning age affects the short- and long-term development of the pig gut microbiome. In this study, we weaned pigs at three different ages, 14, 21, and 28 days, and collected fecal samples 20 times from the neonatal stage until they reached market weight. The fecal microbiome and resistome were assessed using 16S rRNA gene and shotgun metagenomic sequencing to determine how weaning age affected both over the course of the swine production cycle.

## Results

### Effect of weaning age on pig performance

As expected, all pigs gained less weight in the 7 d post-weaning period compared to pigs that were either still nursing or had already been on solid feed for longer than 7 d (Fig. 1). From d 35 onward pigs from all weaning age groups grew at the same rate. There was also no association with weaning age and a pig being removed from the study due to antimicrobial treatment or death (P > 0.05).

**Figure 1.**
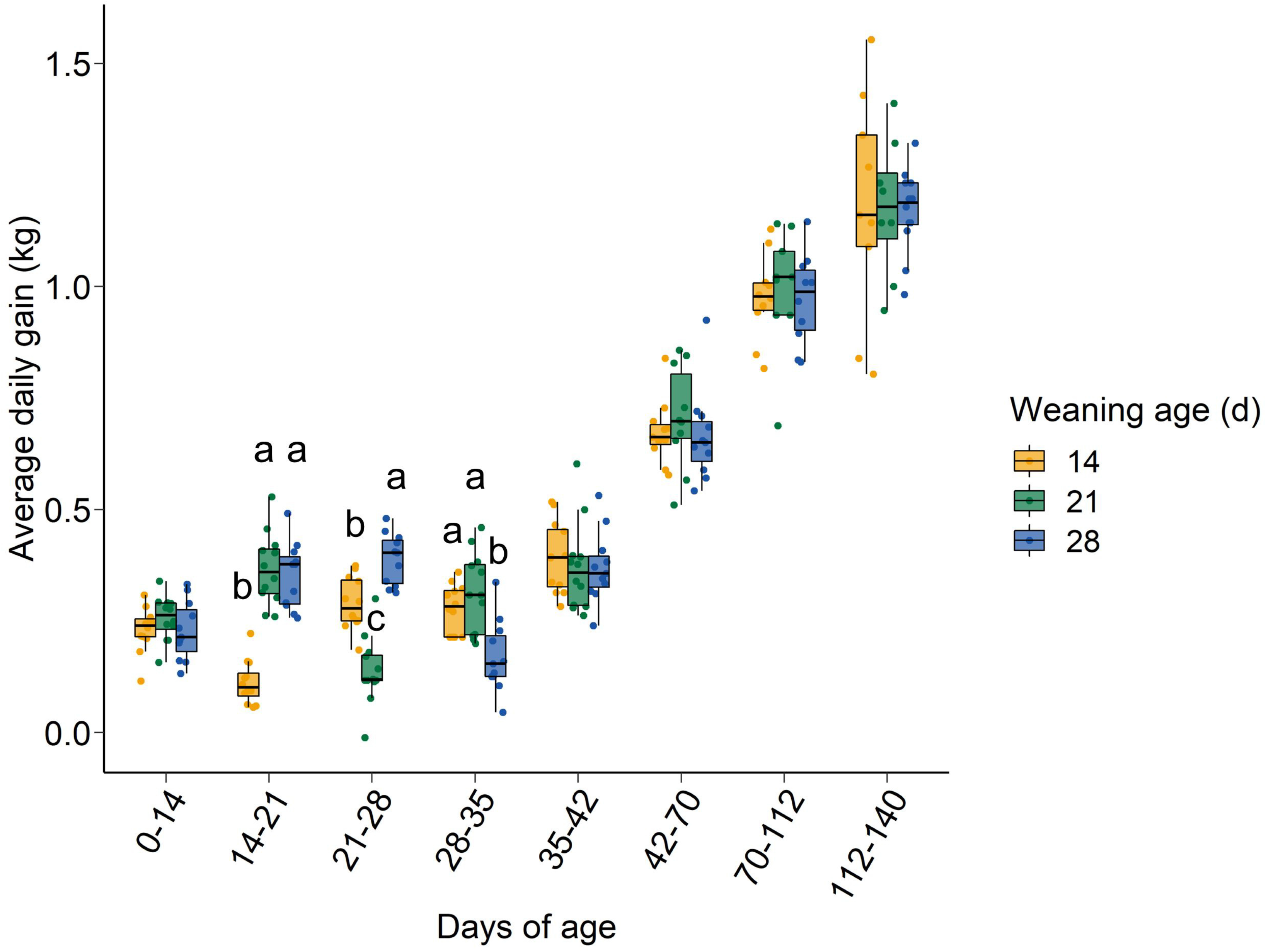
Average daily gain in kg of pigs by weaning age within each weighing period. Different lowercase letters indicate significantly different means (P < 0.05).

### Sequencing

The 16S rRNA gene sequencing of the mock community reflected the expected composition with minor exceptions. There was a larger than expected relative abundance of *Clostridium* (Table S1) and an absence of *Cutibacterium acnes* (formerly *Propionibacterium acnes*); however, this species is known to be poorly amplified by the primers used in this study (11). After processing, there were 35,448 ± 1,247 SEM 16S rRNA gene sequences and 16,699,263 ± 680,292 shotgun metagenomic paired-end sequences per sample. For the metagenomic samples, host contamination accounted for 42.3% ± 2.0% of the sequences.

### Weaning age and the development of the gut microbiome

Weaning age had a strong but temporary effect on the gut microbial community structure (Fig. 2; S2; Fig. S1). Within three days of weaning (d 18), the d 14-weaned pigs had a gut microbiota that was significantly different from that of the pigs that were still nursing (PERMANOVA: R^2^ > 0.25. P < 0.001). By 25 days of age, the gut microbiota of piglets weaned on d 21 was significantly different from that of both the d 14- and d 28-weaned groups (PERMANOVA: R^2^ ≥ 0.13. P < 0.001). However, on d 28, the d 14- and d 21-weaned piglets largely clustered together and separately from the d 28-weaned piglets which were still nursing up to that point. Interestingly, the gut microbial community structure of piglets weaned at 28 days of age remained significantly different from that of the d 14-weaned pigs at d 35 and from the d 21-weaned pigs until and including d 42.

**Figure 2.**
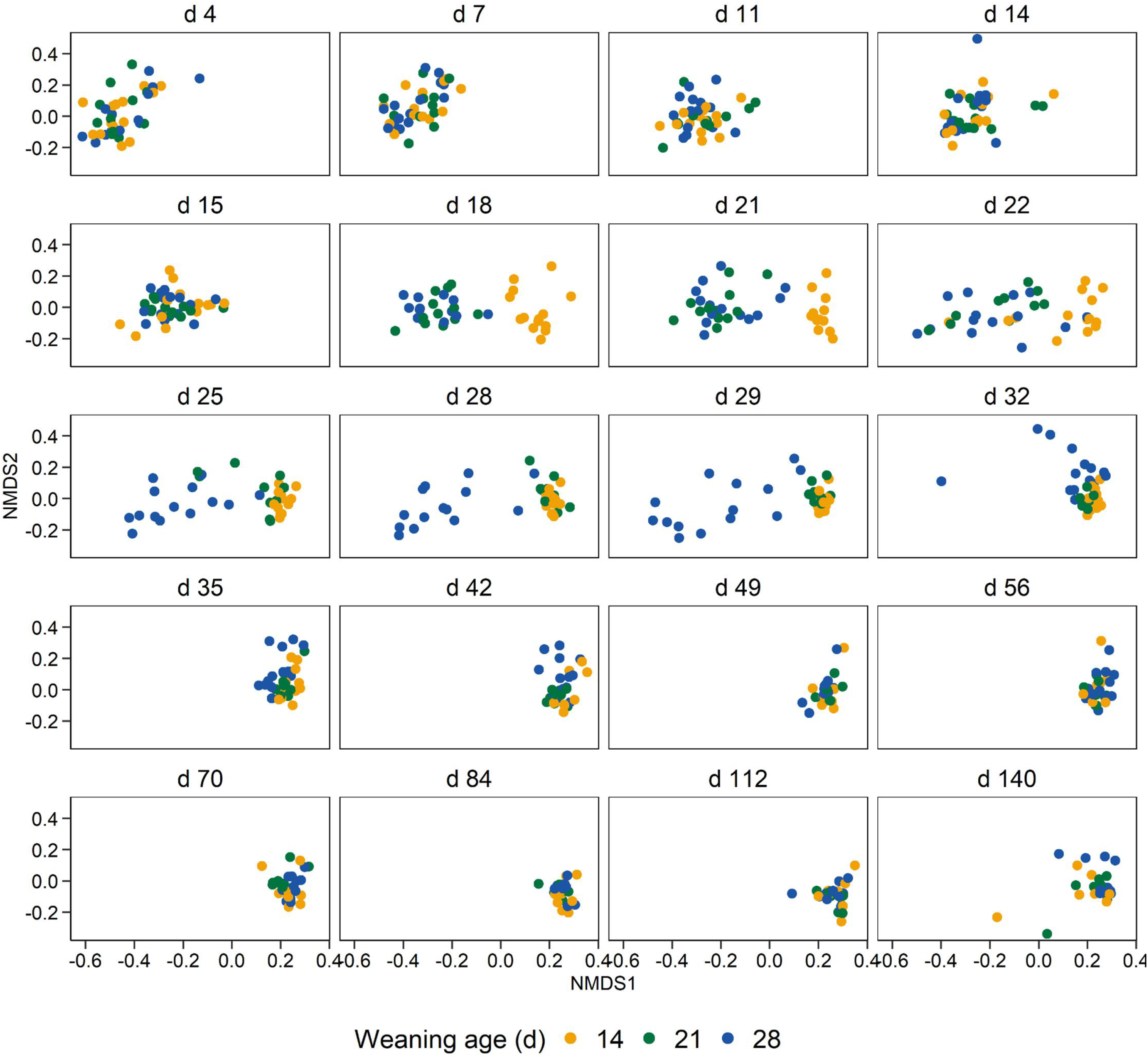
Non-metric multidimensional scaling plot of the Bray-Curtis dissimilarities for the pig fecal microbiota by weaning age and sampling day based on 16S rRNA gene sequencing.

There was an increase in richness (number of OTUs) and diversity (Shannon diversity index) four days post-weaning in the d 14-weaned piglets compared to the still-nursing piglets (Fig. 3A, B). Similarly, from d 25 to 29, both the d 14- and d 21-weaned piglets had greater diversity and richness than the still-nursing d 28-weaned group. These differences had disappeared by d 32, and with the exception of d 42 when the d 28-weaned piglets had a richer microbiota than the other two groups, the diversity of the piglet gut microbiota was not affected by age at weaning. Based on the shotgun metagenomic sequencing analysis, the shifts observed in the gut microbiome post-weaning were associated with a number of different bacterial species (Fig. 3C; see tables S3 and S4 at https://doi.org/10.6084/m9.figshare.c.5619817.v1). Among those that increased in relative abundance post-weaning were several *Prevotella* spp. including *Prevotella copri*, *Prevotella pectinovora*, *Prevotella* sp. P2-180, *Prevotella* sp. P3-122, and *Prevotella stercorea*. *Butyricicoccus porcorum*, *Faecalibacterium prausnitzii*, *Selenomonas bovis*, and *Treponema porcinum* were also among those significantly enriched in pigs that had been weaned at either d 14 or 21 compared to piglets that were not weaned until d 28 (P < 0.05).

**Figure 3.**
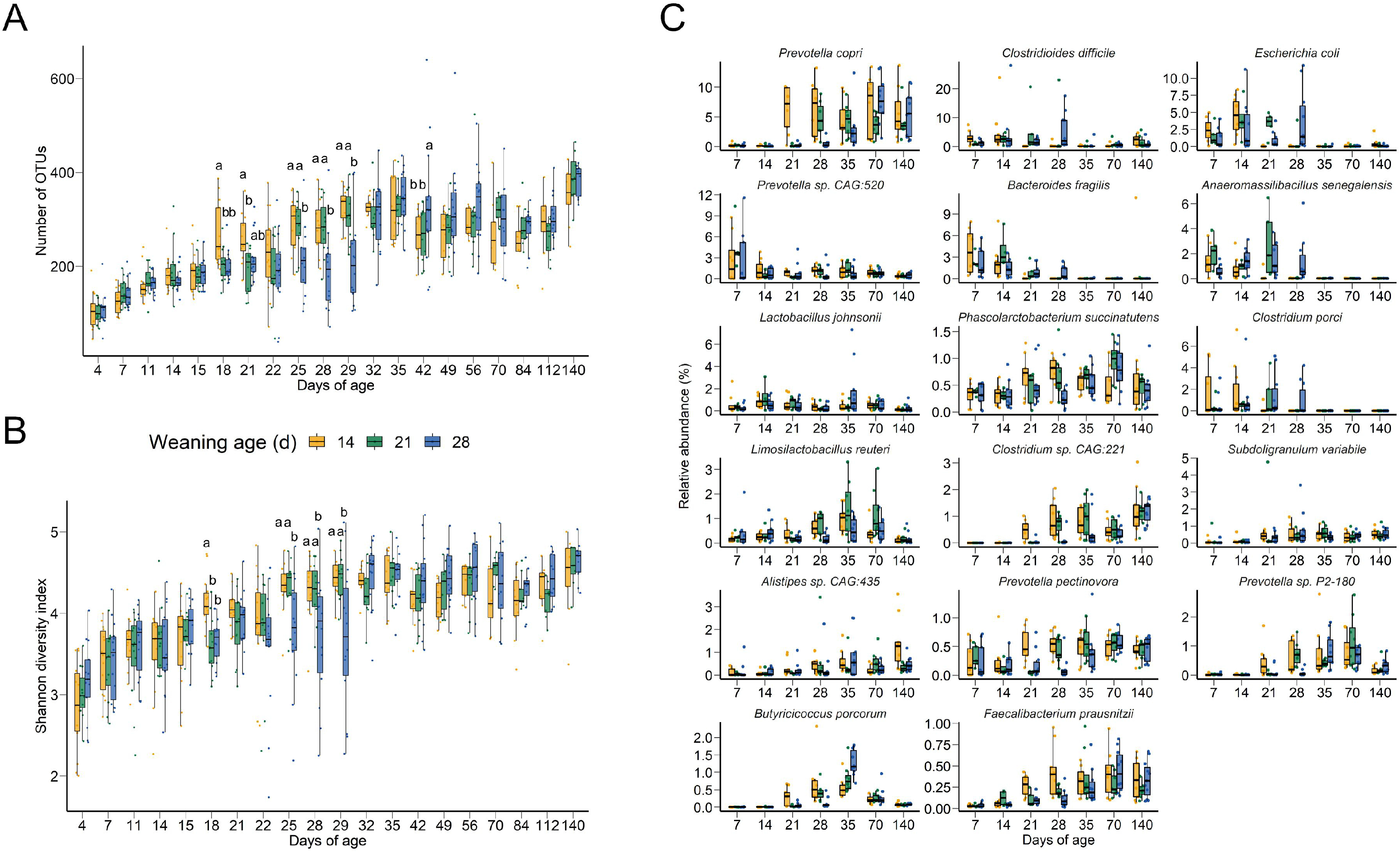
The A) number of OTUs and B) Shannon diversity index values based on 16S rRNA gene sequencing and C) the 15 most relatively-abundant bacterial species based on shotgun metagenomic sequencing for the pig fecal microbiome by weaning age and sampling day. In A and B, different lowercase letters indicate significantly different means (P < 0.05). In C, species are ordered by overall percent relative abundance. *Butyricicoccus porcorum* and *Faecalibacterium prausnitzii* are also included based on their enrichment post-weaning and butyrate-producing activities.

Bacterial species that were consistently associated with nursing pigs included *Anaeromassilibacillus senegalensis*, *Bacteroides fragilis*, *Clostridioides difficile*, *Clostridium porci*, *Clostridium scindens*, *Desulfovibrio piger*, *Escherichia coli*, *Phocaeicola vulgatus*, and *Shigella sonnei* (Table S4). At 35 days of age, only three bacterial species were differentially relatively abundant between the d 14- and d 21-weaned pigs and those weaned on d 28: *Bariatricus massiliensis*, *B. porcorum*, and *D. piger*, all of which were enriched in the d 14- weaned pigs (Table S4). Once the pigs had reached 70 days of age, there were no bacterial species with a relative abundance greater than 0.1% that differed among the groups (P > 0.05).

### Functional changes in the microbiome post-weaning

Functional profiling of the gut microbiome was carried out using the MetaCyc metabolic pathway database and the CAZy database of carbohydrate-active enzymes (CAZymes). The relative abundance of the CAZymes and MetaCyc pathways shifted in a similar way to the microbial taxa post-weaning (Fig. 4A, B). The CAZymes are grouped into the following classes: auxiliary activities (AAs), carbohydrate esterases (CEs), glycoside hydrolases (GHs), glycosyltransferases (GTs), polysaccharide lyases (PLs), and carbohydrate-binding modules (CBMs) which have no enzymatic activity but aid and enhance the catalytic activity of other CAZymes. In total, 237 CAZy families were detected among all samples (Table S5) in comparison with only 61 found within the pig genome (Table S6). All of the CAZyme classes decreased in relative abundance after weaning (Fig. 4C). Overall, 61.5% of the CAZymes were classified as glycoside hydrolases and 24.7% as glycosyltransferases. However, there were still a number of CAZy families that were enriched in the gut microbiomes of post-weaned pigs compared to those still nursing (Table S7). The only AA identified was AA10 (copper-dependent lytic polysaccharide monooxygenases) and in only 35 of the samples (Table S5).

**Figure 4.**
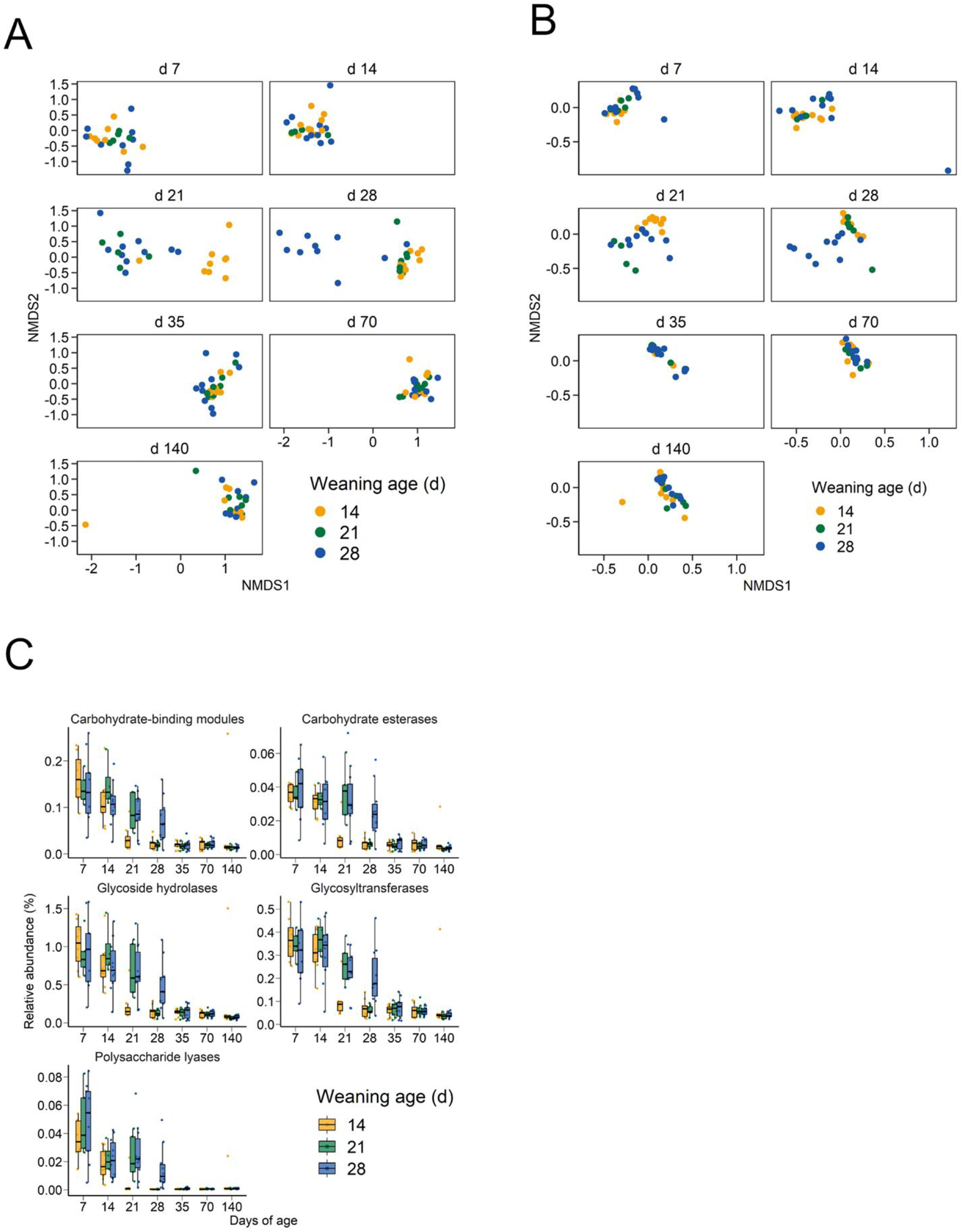
Non-metric multidimensional scaling plot of the Bray-Curtis dissimilarities for the A) CAZymes and B) MetaCyc metabolic pathways of the pig fecal microbiome and C) percent relative abundance of CAZyme classes by weaning age and sampling day.

At 21 days of age, there were 141 unique CAZy families that were differentially abundant between the d 14-weaned pigs and the d 21- and 28-weaned piglets that were still nursing (P < 0.05; Table S7). Similarly, at 28 days of age, 134 CAZy families were differentially abundant between the still nursing d 28-weaned piglets and the post-weaned d 14- and d 21-weaned pigs (p < 0.05; Table S7). There were no differences in CAZy family relative abundance among the three weaning age groups by d 35 (P > 0.05). Many of the alterations in the CAZyme profiles post-weaning reflect the change in diet with CAZy families with lactose-degradation activity (GH2 and GH42) and activity against other components of porcine milk oligosaccharides (PMOs) (GH16, GH18, GH20, GH29, GH30, GH35, GH95, GH139, and GH141) enriched in pigs that were nursing compared to those that had been weaned. Meanwhile, CAZy families including CBMs with mannan- pectin-, starch-, and xylan-binding functions (CBM23, CBM25 CBM26, and CBM77) and GHs with activity against plant cell carbohydrates (GH5, GH39, GH48, GH53, GH93, and GH94) (12, 13), were more relatively abundant in post-weaned pigs that were consuming only a plant-based solid feed.

A large number of MetaCyc metabolic pathways were also differentially abundant between weaned and nursing piglets at d 21 (196 unique pathways) and d 28 (231 unique pathways) with the majority enriched in the gut microbiome of nursing piglets (see Tables S8 and S9 at https://doi.org/10.6084/m9.figshare.c.5619817.v1). As with the CAZymes there was an enrichment of MetaCyc pathways involved in fucose and lactose degradation in the nursing piglets and an increased relative abundance of certain starch degradation pathways post-weaning.

### Weaning age and the gut resistome

Antimicrobial resistance remains a serious challenge to the swine industry and therefore we also characterized the antimicrobial resistome of the pigs longitudinally and in response to weaning age. Similar to the functional analysis, samples clustered by weaning age on d 21 and d 28 when assessed using the relative abundance of antimicrobial resistance genes (ARGs) (Fig. 5A). The large majority of ARGs that were differentially abundant were enriched in the nursing piglets compared to the weaned pigs (see Table S10 at https://doi.org/10.6084/m9.figshare.c.5619817.v1). Notable ARGs that were more relatively abundant in the weaned pigs included *bla*_ACI-1_, *cfxA6*, *erm*(Q), *tet*(44), and *tet*(L). The relative abundance of ARGs conferring resistance to multiple drugs, aminoglycosides, polypeptides, and quinolones as well as several other drug classes decreased post-weaning in all weaning age groups (Fig. 5B). However, tetracycline resistance genes remained relatively stable throughout the pig production cycle. Of the 250 unique ARGs detected, *tet*(Q), *tet*(W), *tet*(O), *aph(3′)-IIIa, mel*, *tet*(W/N/W), *tet*(40), and *tet*(44) were the most relatively abundant among all samples (see Table S11 at https://doi.org/10.6084/m9.figshare.c.5619817.v1).

**Figure 5.**
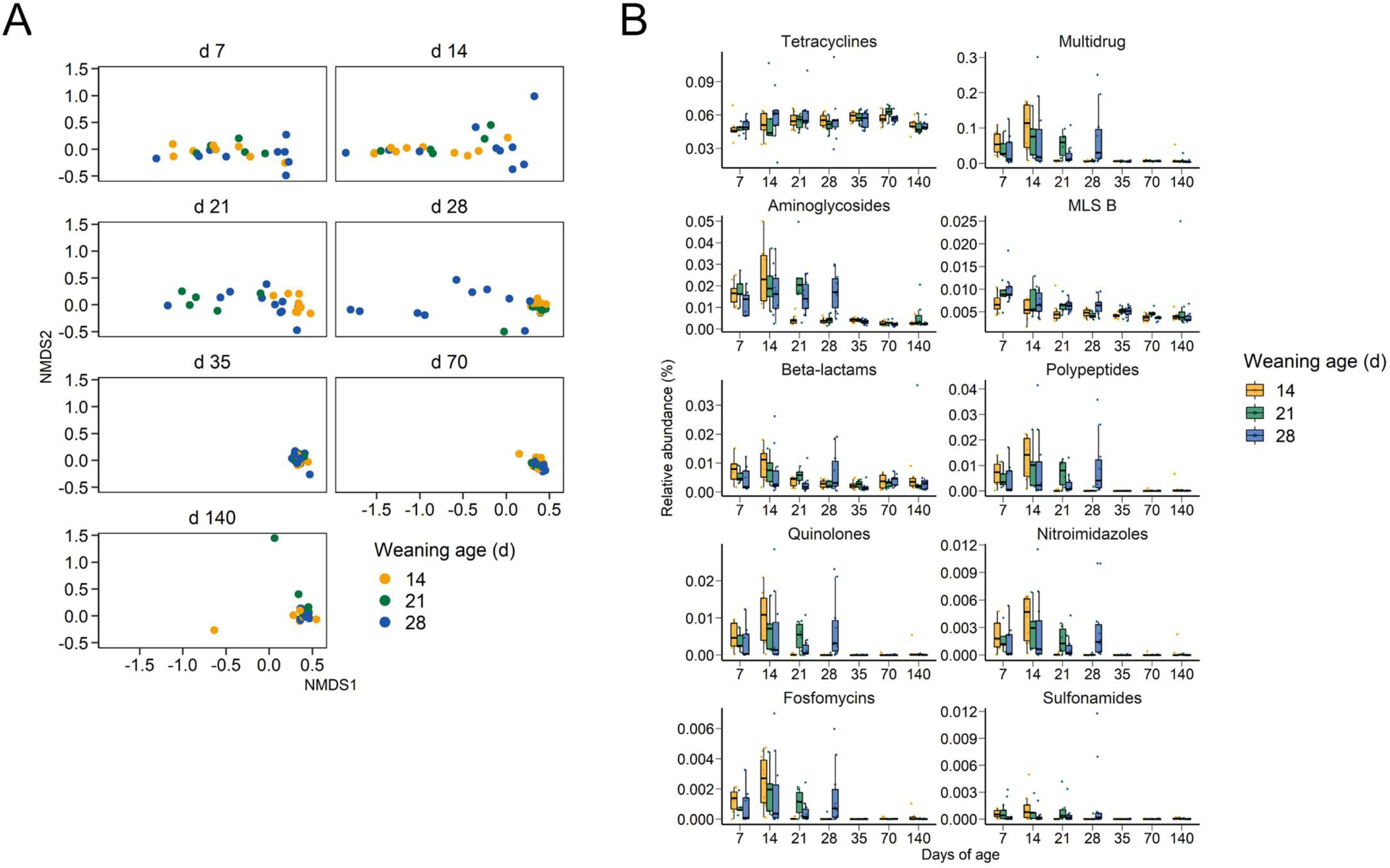
Non-metric multidimensional scaling plot of the Bray-Curtis dissimilarities for the A) antimicrobial resistance genes and B) percent relative abundance of antimicrobial resistance genes by antimicrobial class by weaning age and sampling day.

## Discussion

As expected, there was a substantial shift in the pig gut microbiome within three days of weaning. The sudden change from a milk-based diet to one that is plant-based and less digestible by the pig is largely responsible for this shift immediately post-weaning (1, 14). However, weaning age had no apparent long-term effects on the gut microbiome or the average daily gain of the pigs. A recent study by Massacci et al. (15) that also weaned pigs at different ages (14, 21, 28, and 42 d) with sampling up to 60 d of age also reported no weaning age effect on the microbial community structure at 60 d. Therefore, it appears that a later weaning age only delays post-weaning changes in the gut microbiome rather than affecting the assembly and stability of the microbial community.

Several short-chain fatty acid-producing bacterial species were prevalent among those that were more relatively abundant in pigs that had been weaned. These included *Anaerovibrio slackiae* (acetate, propionate), *B. porcorum* (butyrate), *Coprococcus catus* (butyrate, propionate), *F*. *prausnitzii* (butyrate), *Megasphaera elsdenii* (acetate, butyrate, propionate), *Phascolarctobacterium succinatutens* (propionate), *P*. *copri* (acetate), *Prevotella mizrahii* (acetate), *P*. *pectinovora* (acetate), and *S*. *bovis* (acetate, propionate) (16–21). Short-chain fatty acid-production occurs mostly in the lower gastrointestinal tract of pigs as a result of bacterial fermentation of undigested carbohydrates (22). Acetate, butyrate, and propionate have anti-inflammatory effects on the host (23) and provide up to 25% of daily energy requirements in pigs (24). Butyrate in particular is the primary energy source of colonocytes and regulates apoptosis (25).

Interestingly, *F*. *prausnitzii* has also been reported to be more relatively abundant at 60 days of age in pigs weaned at 21, 28, and 42 d vs. 14 d (15) and in healthy pigs vs. those with post-weaning diarrhea (26). *Butyricicoccus porcorum* has been associated with higher feed efficiency as have *Treponema porcinum* and *Treponema succinifaciens* which were also more relatively abundant in weaned pigs here (27). In-feed supplementation with *Butyricicoccus pullicaecorum* has been shown to improve health and feed efficiency in broiler chickens (28) and *F*. *prausnitzii* reduced intestinal permeability and cytokine expression in a mouse colitis model (29). Therefore, *B*. *porcorum* and *F*. *prausnitzii*, as well as potentially other bacterial species that were more relatively abundant in weaned pigs, are attractive targets for microbiome manipulation and further study into their role in pig gut health.

The bacterial species that were enriched in the microbial communities of pigs that were still nursing at d 21 and d 28 include several potentially pathogenic species such as *C*. *difficile*, *E*. *coli*, *S*. *sonnei*, and *Streptococcus suis*. It is difficult to assess virulence of these species here, however, the presence of potentially pathogenic bacteria pre-weaning may be a risk factor for post-weaning morbidity and mortality (30). Although, many of the more relatively abundant bacterial species were differentially abundant pre- and post-weaning, several remained relatively stable throughout the pig production cycle. In particular, *Lactobacillus johnsonii*, *Mogibacterium kristiansenii*, and *Subdoligranulum variabile* were not affected by weaning. Among these bacterial species *L*. *johnsonii* is the best described and has been reported to improve sow reproductive performance (31) and average daily gain in piglets during the first 35 days of life (32) when delivered in feed. *S*. *variabile*, a butyrate-producing bacterial species, is the only member of its genus and has been previously reported to be a member of the “core microbiota” of the pig gastrointestinal tract (33). *M. kristiansenii* has only recently been described and was originally isolated from pig feces (18).

The functional profile of the gut microbiome also shifted after weaning in all weaning age groups similar to that of the taxonomic profiles. This included a decrease in the relative abundance of all CAZy families post-weaning. The CAZymes encoded by the pig genome are greatly outnumbered by those in the gut microbiome thereby providing the host with an additional source of energy as discussed earlier. Sow’s milk contains not only lactose but at least 119 PMOs (34) which are composed of the monosaccharides fucose, galactose, glucose, N-acetylglucosamine, N-acetylgalactosamine, and sialic acid bound to a lactose or N-actelyllactosamine core (35). These PMOs are generally resistant to host digestive enzymes in the small intestine and are instead fermented by the colonic microbiome into SCFAs (36, 37).

In humans, *Bifidobacterium* and *Bacteroides* spp., including *B*. *fragilis and P. vulgatus* (formerly *Bacteroides vulgatus*), have been shown to metabolize human milk oligosaccharides (38). *Bacteroides fragilis*, which was among the most relatively abundant species in nursing piglets here, carries a number of glycoside hydrolase family genes that facilitate breakdown of milk oligosaccharides (39). Metabolites from the degradation of sialylated bovine milk oligosaccharides by *B*. *fragilis* has also been shown to enhance the growth of *E*. *coli* in vitro (40). All of the GH families found in *B*. *fragilis*, i.e., GH2, GH16, GH18, GH20, GH29, GH33, and GH95, were enriched in the gut microbiomes of nursing piglets. Similarly, GH families and CBMs associated with degradation of plant polysaccharides were more relatively abundant in fecal samples from pigs that had been weaned and consuming a solid plant-based diet for at least 7 d.

The relative abundance of ARGs within several antimicrobial classes decreased post-weaning. However, ARGs conferring resistance to the tetracycline and MLSB classes remained relatively stable throughout the study despite the fact that none of the pigs were exposed to any antimicrobials. Not surprisingly, these are the antimicrobial classes with the longest history of use in swine production and are still among the most frequently administered antimicrobials in North American pigs (41, 42). This background level of tetracycline and MLSB resistance probably also explains why several studies have reported limited or only temporary effects on the pig gut microbiome following exposure to drugs of these antimicrobial classes (8, 43, 44). The reason for the significant decrease in other ARGs after weaning is likely due to the post-weaning shift in bacterial taxa carrying these ARGs. For example, many of the relatively abundant multidrug ARGs such as *mdtF*, *acrF*, *evgS*, *acrB*, *mdtO*, *mdtP*, and *cpxA*, are found in the majority of *E. coli* and *S*. *sonnei* genomes and both of these species decreased in relative abundance post-weaning. In contrast, relatively abundant tetracycline resistance genes such as *tet*(Q), *tet*(W), and *tet*(O) have a much wider host range (45).

Two of the ARGs that were more relatively abundant in weaned piglets compared to those still nursing were the Ambler class A beta-lactamase genes bla_CfxA6_ and *bla*_ACI-1_. Additionally, bla_CfxA2_ was enriched in piglets weaned at d 14 and 21 compared to those still nursing on d 28. Both bla_CfxA2_ and bla_CfxA6_ have been identified in several *Prevotella* spp. (46) which likely accounts for the post-weaning enrichment of these ARGs. In *Prevotella* spp. the *bla*_CfxA_ genes have been shown to confer resistance to ampicillin but not cefmetazole (47). The *bla*_ACI-1_ gene may be associated with *M*. *elsdenii* as has been demonstrated in human gut metagenomes (48). Overall, these results again demonstrate the challenges faced when it comes to reducing antimicrobial resistance in swine as none of the pigs in this study were exposed to antimicrobials.

In conclusion, this study shows that weaning age has little effect on the long-term development and composition of the pig gut microbiome and resistome. Instead, the pig gut microbiome tends to change in a rather predictable manner post-weaning in a swine production environment. Many ARGs also persisted in the feces of the pigs throughout the study likely reflecting the long history of use of certain antimicrobial classes in swine production. Several bacterial species with potential beneficial properties such as SCFA production were found to be enriched post-weaning and are attractive targets for future microbiome manipulation and culture-based studies.

## Experimental Procedures

### Animals and experimental design

All pig experiments were carried out at the swine unit of the Lacombe Research and Development Centre. Seven pregnant sows that farrowed within 24 h of each other were used in the study. A total of 45 piglets (n = 15 per weaning age group) were randomly selected for inclusion in the study based on litter, weight, and sex, with low-weight piglets excluded. Following weaning, all pigs were fed the same starter diet that was free of antibiotics, prebiotics, and probiotics (see Table S12 at https://doi.org/10.6084/m9.figshare.c.5619817.v1). Any pig that required an antibiotic treatment was removed from the study. Animals in this experiment were cared for in agreement with the Canadian Council for Animal Care (2009) guidelines. The Lacombe Research and Development Centre Animal Care Committee reviewed and approved all procedures and protocols involving animals.

On d 4 prior to sampling, 15 piglets were randomly chosen from among the 7 litters and designated to be weaned at 14, 21, or 28 days of age (Fig. S2). Piglets were sampled using fecal swabs (FLOQSwabs, Copan, Murrieta, CA, USA) beginning at 4 days of age and repeated on days 7 and 11. At 14 days of age, piglets assigned to the d 14 weaning group were removed from their sow after sampling and transferred to a nursery room within the swine barn. Fecal sampling continued for all piglets at 15, 18, and 21 d of age. On d 21, piglets in the d 21 weaned group were removed from their sow and placed in a nursery room. Fecal samples were taken from all pigs on d 22, 25, and 28, and on d 28 the d 28-weaned piglets were weaned from their sow and placed in the nursery room. Piglets were then sampled on d 29, 32, 35, 42, 49, 56, 70, 84, 112, and 140. All fecal swabs were immediately placed on ice, transported to the laboratory, and stored at −80°C until DNA extraction.

### DNA extraction and 16S rRNA gene and shotgun metagenomic sequencing

DNA was extracted from fecal material collected on FLOQSwabs with the QIAamp BiOstic Bacteremia DNA Kit (Qiagen, Mississauga, ON, Canada) as per manufacturer’s instructions with the following modifications. Sterile scissors were used to remove the swab which was then placed into a PowerBead tube with MBL solution and agitated at 70°C and 400 rpm for 15 min. After heating, the tubes were shaken in a FastPrep-24 (MP Biomedicals, Solon, OH, USA) at 4.0 m/s for 45 s. Tubes were allowed to rest in the MP FastPrep-24 for 5 min. Using sterile forceps, swabs were removed from PowerBead tubes prior to pelleting debris at 10,000 x g for 2 min. All remaining steps were followed as per the manufacturer’s protocol.

Extracted bacterial DNA was loaded onto nine 96-well plates and two wells on each plate included a positive control (MSA-1002, 20 Strain Even Mix Genomic Material, ATCC, Manassas, VA, USA) and negative control (water). Negative extraction controls were also included. DNA was quantified and analyzed using the Qubit dsDNA HS Assay Kit (Thermo Fisher Scientific, Waltham, MA, USA) and Agilent High Sensitivity D1000 ScreenTape System (Santa Clara, CA, USA). The V4 hypervariable region of the 16S rRNA gene was amplified as per Kozich et al. (49). To prepare each 16S rRNA gene library, 5 μl of each sample from three 96-well plates were pooled at a time. The pooled library was normalized to 0.4 nM and submitted to the Genomics Facility in the Infectious Bacterial Diseases Research Unit at USDA-ARS-NADC in Ames, IA for 250 bp paired-end sequencing on a MiSeq instrument (Illumina, San Diego, CA) using v2 chemistry.

DNA from d 7, 14, 21, 28, 35, 70, and 140 of all pigs that remained in the study through to d 140 was also subjected to shotgun metagenomic sequencing. Metagenomic libraries were prepared using 700 ng of DNA and the TruSeq DNA PCR-Free Library Prep Kit (Illumina Inc.) following the manufacturer’s recommended protocol. Briefly, DNA was fragmented to an average length of 400 bp with a Covaris LE220 instrument, end-repaired, A-tailed, and indexed with TruSeq Illumina adapters. Libraries were then validated on a Fragment Analyzer system with a High Sensitivity NGS Fragment Kit (Agilent Technologies, Mississauga, ON, Canada) to check for size and quantified by qPCR using the Kapa Library Quantification Illumina/ABI Prism Kit protocol (KAPA Biosystems, Wilmington, MA, USA). Equimolar quantities of each library were then pooled and sequenced on the Illumina NovaSeq 6000 instrument with a SP flowcell (2 x 250 bp) following manufacturer’s instructions.

### 16S rRNA gene sequence analysis

The 16S rRNA were processed using DADA2 v. 1.14 (50) in R v. 3.6.3. Briefly, the forward and reverse reads were trimmed to 200 and 210 bp, respectively, merged with a minimum overlap of 75 bp, and chimeras removed. The RDP naive Bayesian classifier (51) and the SILVA SSU database release 138 (52) were then used to assign taxonomy to each merged sequence, referred to here as operational taxonomic units (OTUs) with 100% similarity. OTUs that were classified as chloroplasts, mitochondria, or eukaryotic in origin, and those that were identified in the extraction control samples at an equal or higher abundance than the biological samples were removed prior to analyses. The number of OTUs, Shannon diversity index, inverse Simpson’s diversity index, and the Bray-Curtis dissimilarities were calculated in R v. 4.0.0 using Phyloseq 1.32.0 (53) and vegan v. 2-5.6 (54). To account for uneven sequencing depth all samples were randomly subsampled to 6,900 sequences per sample prior to analyses.

### Metagenomic sequence analysis

Metagenomic sequences were trimmed (quality score < 15 over a sliding window of 4 bp; minimum length of 50 bp) and sequencing adapters removed using Trimmomatic v. 0.38 (55). Bowtie2 v. 2.4.2-1 (56) was used to align host sequences to the *Sus scrofa* genome (Sscrofa11.1) for removal. Taxonomy was assigned to the filtered metagenomic sequences using Kaiju v. 1.7.3 (57) and the NCBI non-redundant protein database (October 13, 2020). For functional profiling of the metagenomic samples, HUMAnN v. 3.0.0.alpha.1 (58) was used to align reads to the UniRef90 database which were then collapsed into MetaCyc metabolic and enzyme pathways (59). Reads were aligned to the Comprehensive Antibiotic Resistance Database (CARD) v. 3.0.8 (60) and the Carbohydrate-Active enZYmes (CAZy) Database (dbCAN2) v. 07312020 (61) using DIAMOND v. 0.9.28 (62) (≥ 90% amino acid identify and ≥ 90% coverage).

### Statistical analysis

Fisher’s exact test was used to determine if weaning age was associated with removal from the study post-weaning due to antimicrobial treatment or death. The effect of weaning age on the microbial community structure was assessed using the Bray-Curtis dissimilarities and PERMANOVA (adonis2 function). The R package pairwiseAdonis (63) was used to compare the Bray-Curtis dissimilarities within each sampling time and the Benjamini-Hochberg procedure was used to correct P-values for multiple comparisons. The effect of weaning age on the relative abundance of microbial species, CAZy families, MetaCyc pathways, and ARGs was determined using MaAsLin2 (microbiome multivariable associations with linear models) v. 1.5.1 (64) in R. Only those microbial species with an average relative abundance of at least 0.1% and CAZy families, MetaCyc pathways, and ARGs identified in at least 25% of samples were included in these analyses.

### Data Availability

All 16S rRNA gene and metagenomic sequencing data are available at the NCBI sequence read archive under BioProject PRJNA629856.

## Acknowledgements

We thank the animal care staff at the Lacombe Research and Development Centre swine unit for their assistance and excellent care of the animals. We are also grateful to Cara Service and TingTing Liu for their valuable technical support. We thank David Alt and Crystal Loving for help with sequencing and Arun Kommadath for submitting the metagenomic sequences to the SRA. We also appreciate helpful feedback from Cassidy Klima. This research was supported by Alberta Agriculture and Forestry grant 2018R009R and also used resources provided by the SciNet project of the USDA ARS, ARS project number 0500-00093-001-00-D.

## Supplemental Material

**Figure S1.** Non-metric multidimensional scaling plot of the Bray-Curtis dissimilarities for the fecal microbiota by weaning age and age of piglets.

**Figure S2.** Frequency of fecal sampling of the pigs in this study.

**Table S1.** Percent relative abundance of genera identified in the mock communities (ATCC MSA-1002) via 16S rRNA gene sequencing. Note: *Schaalia* spp. are classified as *Actinomyces* spp. within the SILVA SSU database.

**Table S2.** Pairwise PERMANOVA of the Bray-Curtis dissimilarities by weaning age within each sampling time.

**Table S3.** Percent relative abundance (± SEM) of microbial species by weaning age and sampling time based on metagenomic sequencing. Only those microbial species with a relative abundance greater than 0.01% are included. Species are listed by overall percent relative abundance.

**Table S4.** Differentially abundant microbial species based on metagenomic sequencing. Negative coefficient values (highlighted) at 21 d of age indicate that the microbial species was more relatively abundant in the d 14-weaned pigs vs. the d 21- and 28-weaned pigs. Negative coefficient values (not highlighted) at 28 and 35 d of age indicate that the microbial species was more relatively abundant in the d 28-weaned pigs vs. the d 14- and 21-weaned pigs.

**Table S5**. Percent relative abundance (± SEM) of CAZy families detected in at least one sample by weaning age and sampling time.

**Table S6**. CAZy families detected in the *Sus scrofa* genome at 90% identity.

**Table S7**. Differentially abundant CAZy families on d 21 and 28 between weaned vs. nursing piglets. Negative coefficient values (highlighted) at 21 d of age indicate that the CAZy family was more relatively abundant in the d 14-weaned pigs vs. the d 21- and 28-weaned pigs. Negative coefficient values (not highlighted) at 28 d of age indicate that the CAZy family was more relatively abundant in the d 28-weaned pigs vs. the d 14- and 21-weaned pigs.

**Table S8.** Percent relative abundance (± SEM) of MetaCyc pathways detected in at least one metagenomic sample by weaning age and sampling time.

