## Supplementary figures and images for "Weaning age and its effect on the development of the swine gut microbiome and resistome"

### Supplementary Fig. S1

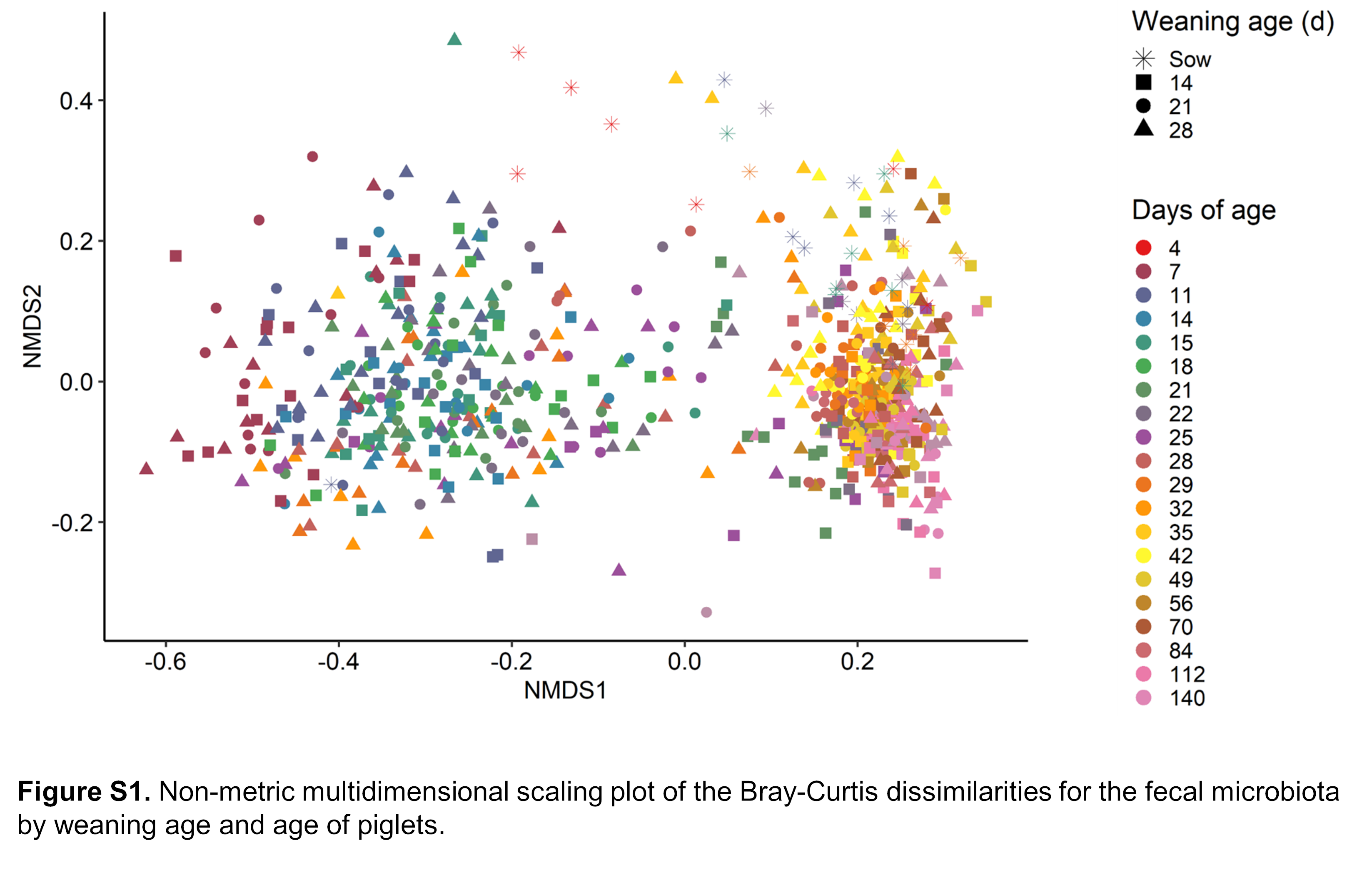

### Supplementary Fig. S2

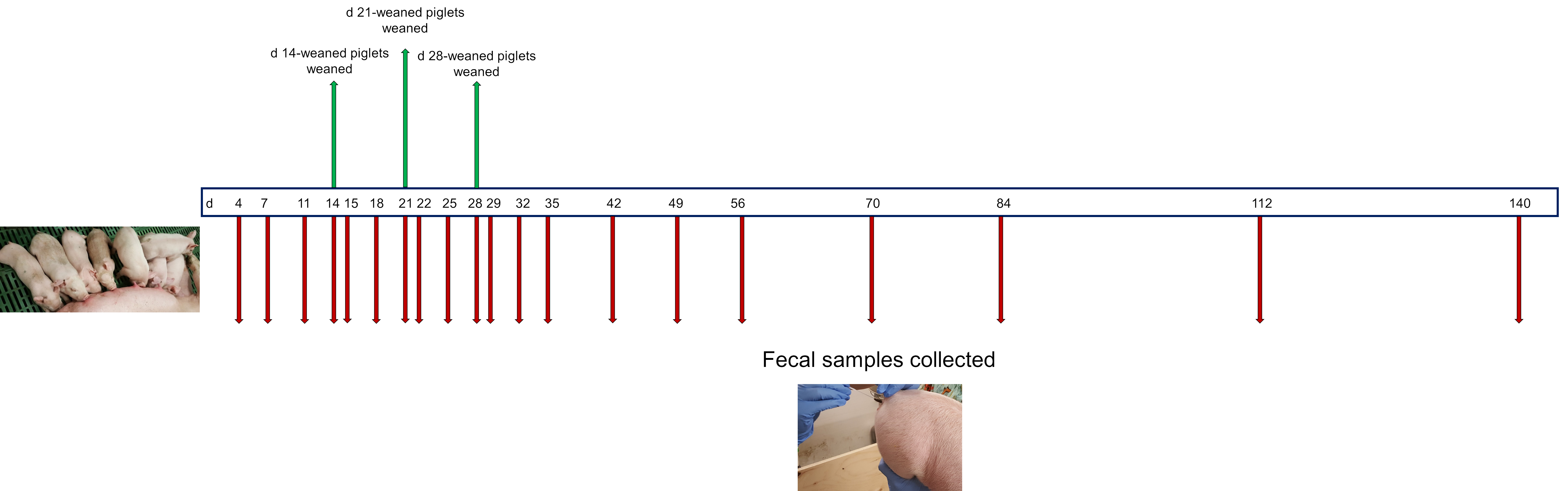
